# Variability in the relationship between ocean phytoplankton diversity and carbon biomass across methods and scales

**DOI:** 10.64898/2026.02.19.706895

**Authors:** Sasha J. Kramer

## Abstract

Diverse ecosystems on land and in the ocean are thought to be more productive, stable, and resistant to change, but these relationships are highly variable across systems and scales. Although the productivity-diversity relationship (PDR) has been extensively explored on land, there are limited in-water observations of the PDR in marine ecosystems. In this work, the relationship between phytoplankton diversity and carbon biomass (as a proxy for productivity) was examined using a global in situ dataset. The shape of this relationship was evaluated for three metrics of phytoplankton diversity: pigment concentrations modeled from hyperspectral remote sensing reflectance (*R_rs_*(*λ*)), pigment concentrations measured by high performance liquid chromatography (HPLC), and 18S rRNA gene sequences (eukaryotes). While gene sequencing methods provide higher resolution taxonomic information about phytoplankton communities, remote sensing methods collect higher resolution spatiotemporal information on global scales. By comparing these methods (*R_rs_*(*λ*)-modeled pigments vs. measured HPLC pigments vs. 18S rRNA gene sequences), this work demonstrates the variability in the relationship between phytoplankton diversity and carbon biomass based on the method of assessing these parameters, and establishes a baseline for in situ observations that has the potential to be extended to global observations from NASA’s Plankton Aerosol Cloud ocean Ecosystem (PACE) satellite.

## 1. Introduction

The relationship between biodiversity and ecosystem function is highly variable and has long been a subject of interest for ecologists across marine and terrestrial environments (Rosenzweig 1995; Chapin III et al. 1997; Longhurst 1998; Mittelbach et al. 2001; Gagné et al. 2020; Sommeria-Klein et al. 2021). Biodiversity is intrinsically linked to ecosystem structure and function (Chapin III et al. 1997). The amount of primary production in an ecosystem supports essential ecosystem services that benefit human health and economies: the continuity of these services depends in part on maintaining high levels of biodiversity, which is a major goal for conservation in the ocean and on land (Pimentel et al. 1992; Worm et al. 2006; Palumbi et al. 2008; Cardinale et al. 2012). More diverse ecosystems have higher functional redundance, characterized by multiple groups of primary producers that contribute similarly to these fundamental ecosystem services (Behrenfeld et al. 2021a; b). Following a major disturbance, ecosystems that are more biodiverse have higher resistance to change and are better able to maintain their pre-disturbance productivity (Baert et al. 2016). Given the importance of biodiversity to ecosystem functioning, the productivity-diversity relationship (PDR) has been extensively studied across scales on land (Chapin III et al. 1997; Mittelbach et al. 2001; Chase and Leibold 2002). However, it is still not well understood for open ocean ecosystems, due in part to the difficulty of surveying stable communities of primary producers in a global, dynamic physical system.

Most primary productivity in the ocean is performed by phytoplankton, microscopic organisms that form the base of the oceanic food web and contribute to biogeochemical cycling and carbon sequestration (Legendre 1990; Field et al. 1998; Arrigo 2005). Phytoplankton are extremely diverse, representing broad functional classes, cell sizes, and taxonomies (de Vargas et al. 2015; Righetti et al. 2019). Marine phytoplankton biogeography is broadly structured by a combination of abiotic factors (light, nutrients, sea surface temperature [SST], physical mixing) and biotic factors (viruses, grazers, predator-prey interactions) (Righetti et al. 2019; Sommeria-Klein et al. 2021). Characterizing the relationship between phytoplankton diversity and productivity is essential for a complete understanding of the Earth system processes that rely on phytoplankton. Furthermore, the correlation between phytoplankton biodiversity and temperature in marine systems suggests that climate change will have significant impacts on the shape of the marine PDR (Ibarbalz et al. 2019; Anderson et al. 2021). As the ocean changes in response to anthropogenic climate change, phytoplankton diversity and productivity will naturally be affected (Behrenfeld 2014; Behrenfeld et al. 2015).

Global Earth system models and laboratory experiments have both considered the impacts of biodiversity change or loss on marine ecosystems (Vallina et al. 2017; Dutkiewicz et al. 2019; Henson et al. 2021). For instance, (Vallina et al. 2014) predicted a unimodal PDR for phytoplankton in the global ocean, similar to that observed for many land plants (Rosenzweig 1995), where diversity is highest at intermediate productivity. However, far fewer studies have actually observed the PDR in situ for marine phytoplankton due, in part, to the difficulty of observing marine systems on the spatiotemporal scales necessary for capturing phytoplankton diversity (Strong et al. 2015; Acevedo-Trejos et al. 2018; Ibarbalz et al. 2019; Gagné et al. 2020). The existing past in situ studies were associated with individual field campaigns (specifically, Tara Oceans (Ibarbalz et al. 2019) and AMT (Acevedo-Trejos et al. 2018)) and did not find the unimodal relationship predicted in global models of the phytoplankton PDR. While it is challenging to compare directly between different proxies for phytoplankton diversity and productivity used in these studies, the broad trends can still be considered. Specifically, (Acevedo-Trejos et al. 2018) observed the highest cell size diversity at high levels of gross primary productivity, while (Ibarbalz et al. 2019) observed an inverse linear relationship between taxonomic diversity and chlorophyll-*a* concentration. Put together, these two studies offer examples of each side of the theorized unimodal PDR curve that is observed on land and predicted by models for the ocean (Vallina et al. 2014): either a monotonically increasing (Acevedo-Trejos et al. 2018) or decreasing (Ibarbalz et al. 2019) relationship between phytoplankton diversity and ecosystem productivity.

In these in situ studies and elsewhere in the literature, phytoplankton carbon biomass, *C_p_*_ℎ*yto*_, is often used as a proxy for productivity, as it is this phytoplankton carbon that will be integrated into the food web or sequestered in sinking particles via the biological pump (Irigoien et al. 2004; Behrenfeld and Boss 2014). Much of the open ocean is near steady state much of the time in terms of biomass (Behrenfeld and Boss 2014), with the exception of episodic disturbances caused by, e.g., storms or predation events, but community composition may be changing rapidly. A combination of bottom-up (e.g., nutrient limitation) and top-down (e.g., grazing) factors can contribute to the relative stability of *C_p_*_ℎ*yto*_ in the subtropical gyres or elevated biomass during the bloom apex at higher latitudes (Behrenfeld 2014; Vallina et al. 2014). The vast diversity of phytoplankton in most systems implies redundancy in the environment across phytoplankton traits (Chen et al. 2019; Behrenfeld et al. 2021a; b), suggesting that an ecosystem characterized by high phytoplankton diversity will be more resistant to environmental change. Modeling studies have demonstrated the importance of characterizing rare disturbances, and of measuring how that disturbance dissipates in the ecosystem over time (Lan Smith et al. 2016). While it is widely recognized that species diversity often does not equal genetic or functional diversity within the phytoplankton, species diversity is still often used as a tool to survey populations (Biller et al. 2015; Li and Chesson 2016). Higher phytoplankton taxonomic diversity is more closely associated with increased *resistance*, or maintaining ecosystem function post-disturbance, rather than increased *resilience*, or full recovery of an identical ecosystem post-disturbance (Baert et al. 2016). At the same time, the dispersed nutrient and prey field in the ocean means that there are fewer individual niches for phytoplankton groups in the ocean than there are for plants on land (Behrenfeld et al. 2021b), so a large ecological disturbance may still impact ecosystem resilience despite a diverse phytoplankton community.

Characterizing the relationship between phytoplankton diversity and carbon biomass will enable us to better monitor and predict responses to a changing Earth system, but this goal will only be achievable by linking observations of phytoplankton diversity and carbon biomass to satellite ocean color (Graff et al. 2015; Kramer et al. 2022; Stramski et al. 2022), which is measured on scales relevant to these global processes. The launch of NASA’s Plankton Aerosol Cloud ocean Ecosystem (PACE) satellite and its Ocean Color Instrument (OCI) (Werdell et al. 2019) in February 2024 has enabled advances in global ocean color with hyperspectral remote sensing reflectance (*R_rs_*(*λ*)). PACE OCI’s hyperspectral resolution allows for improved characterization of global ocean phytoplankton community composition from space (Cetinić et al. 2024), specifically by modeling phytoplankton pigment concentrations from *R_rs_*(*λ*) (Chase et al. 2017; Kramer et al. 2022). The current study aims to take advantage of these new capabilities in ocean color remote sensing to address a knowledge gap in marine microbial ecology by combining datasets that define phytoplankton diversity across similar metrics (specifically, modeled phytoplankton pigments from hyperspectral *R_rs_*(*λ*) data, measured phytoplankton pigments from high performance liquid chromatography [HPLC], and 18S rRNA gene sequences) to characterize the relationship between phytoplankton diversity and carbon biomass on global scales from in situ methods and remote sensing. This work investigates: (1) What is the shape of this relationship for global phytoplankton taxonomic diversity and carbon biomass on regional to global scales? And (2) how does the resolution of phytoplankton taxonomy across methods (from *R_rs_*(*λ*)-modeled pigments, measured HPLC pigments, and 18S rRNA genes) impact the shape of the retrieved relationship?

## 2. Materials and Methods

### 2.1 Global surface ocean dataset

Samples were compiled from publicly available datasets collected in the surface ocean (≤10 meters) for comparison to remote sensing data (**Figure 1**). Five field campaigns collected phytoplankton pigment and 18S rRNA gene sequence data that fit these criteria, for a total of 324 paired surface samples (**Figure S1**). EXport Processes in the Ocean from RemoTe Sensing (EXPORTS; N = 61) included field campaigns in the North Pacific (August-September 2018) and North Atlantic (May 2021) Oceans (Siegel et al. 2021; Johnson et al. 2023). The Malaspina Circumnavigation Expedition (N = 128) sampled the global ocean across tropical to subtropical latitudes from 2010 to 2011 (Duarte 2015). The GEOTRACES GA10 cruise (N = 7) sampled the Southern Atlantic Ocean in October and November 2010 (Wyatt et al. 2021; McNichol et al. 2025b). Tara Oceans and Tara Polar (N = 107) assessed every major ocean basin from 2009 to 2014 (Sunagawa et al. 2020). Finally, the North Atlantic Aerosols and Marine Ecosystems Study (NAAMES; N = 21) occupied the western North Atlantic Ocean once per season from 2015 to 2018 to capture the succession of the phytoplankton bloom (Behrenfeld et al. 2019).

**Figure 1.**
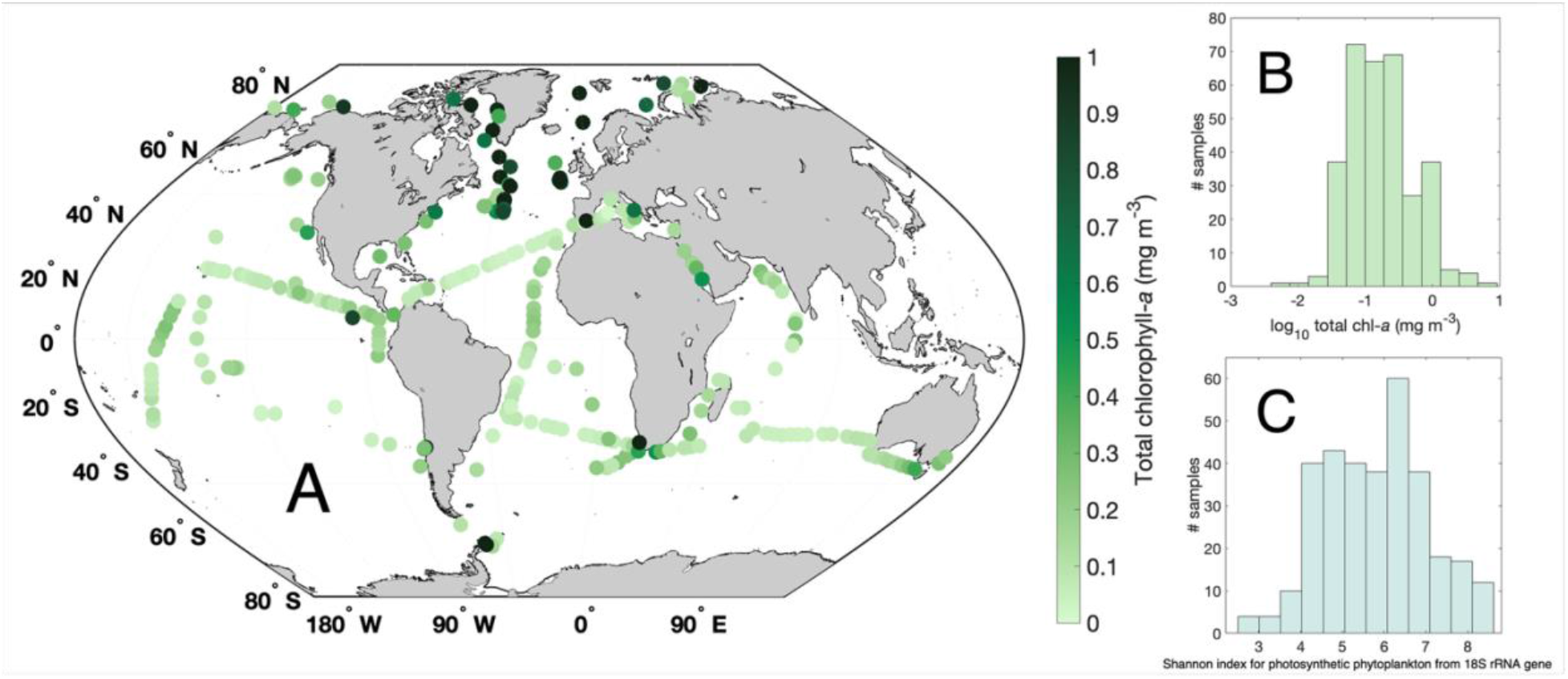
Summary of samples used in this study. (A) Global map of sample distribution, colored by HPLC total chlorophyll-*a* concentration. (B) Histogram of HPLC total chlorophyll-*a* distribution for all samples and (C) histogram of Shannon diversity index for photosynthetic phytoplankton calculated from ASV-level 18S rRNA gene relative sequence abundances.

### 2.2 Phytoplankton pigments modeled from remote sensing reflectance R_rs_(λ)

As a first metric, to assess the phytoplankton diversity that would be visible from PACE, phytoplankton pigment concentrations were modeled from hyperspectral remote sensing reflectance (*R_rs_*(*λ*); sr^-1^) generated by the Spectral Derivatives Pigment (SDP) model (Kramer et al. 2022) for the sample dataset used in that study, with additional paired data from the EXPORTS North Atlantic field campaign as shown in (Kramer et al. 2024b) (N = 162). This sample dataset includes paired HPLC phytoplankton pigment measurements and in situ hyperspectral *R_rs_*(*λ*) measurements. Forty-one samples in that dataset were also included in the present study (from NAAMES, EXPORTS, and Tara Oceans) since they also had paired 18S rRNA gene sequences (Section 2.4). Full details of the SDP model are found in (Kramer et al. 2022) and SDP model code is available on GitHub (see Data Availability Statement). Briefly, the concentrations of thirteen phytoplankton pigments (total chlorophyll-*a* and twelve accessory pigments) are modeled from a residual spectrum calculated between a measured *R_rs_*(*λ*) spectrum and a modeled *R_rs_*(*λ*) spectrum. The twelve pigments retrieved by SDP are: peridinin (Perid), fucoxanthin (Fuco), chlorophyll c1+c2 (Chlc12), chlorophyll c3 (Chlc3), hexanoyloxyfucoxanthin (HexFuco), butanoyloxyfucoxanthin (ButFuco), alloxanthin (Allo), prasinoxanthin (Pras), neoxanthin (Neo), monovinyl chlorophyll b (MVchlb), divinyl chlorophyll a (DVchla), and zeaxanthin (Zea). This model is currently being implemented globally for PACE, so it was applied to in situ data here as a proof-of-concept.

### 2.3 HPLC pigments

The concentrations of total chlorophyll-*a* and accessory pigments were also measured for each field campaign using high performance liquid chromatography (HPLC; **Figure 2**). The sample twelve accessory pigments that are retrieved by SDP were examined from HPLC; these pigments were selected for their taxonomic relevance in describing photosynthetic phytoplankton community composition (Kramer and Siegel 2019; Kramer et al. 2022). Accessory pigment concentrations were normalized to total chlorophyll-*a* concentration to weaken the degree of positive linear correlation with other accessory pigments and total chlorophyll-*a* (Kramer and Siegel 2019). The relative concentration of these phytoplankton pigments to total chlorophyll-*a* was then used to separate five distinct phytoplankton groups using a hierarchical cluster analysis performed based on the correlation distance and Ward’s linkage method (**Figure 2**): dinoflagellates, diatoms, green algae, prymnesiophytes, and cyanobacteria (Jeffrey et al. 2011; Kramer and Siegel 2019). Total chlorophyll-*a* concentrations ranged from 0.006-7.2 mg m^-3^ and global values were normally distributed (**Figure 1B**).

**Figure 2.**
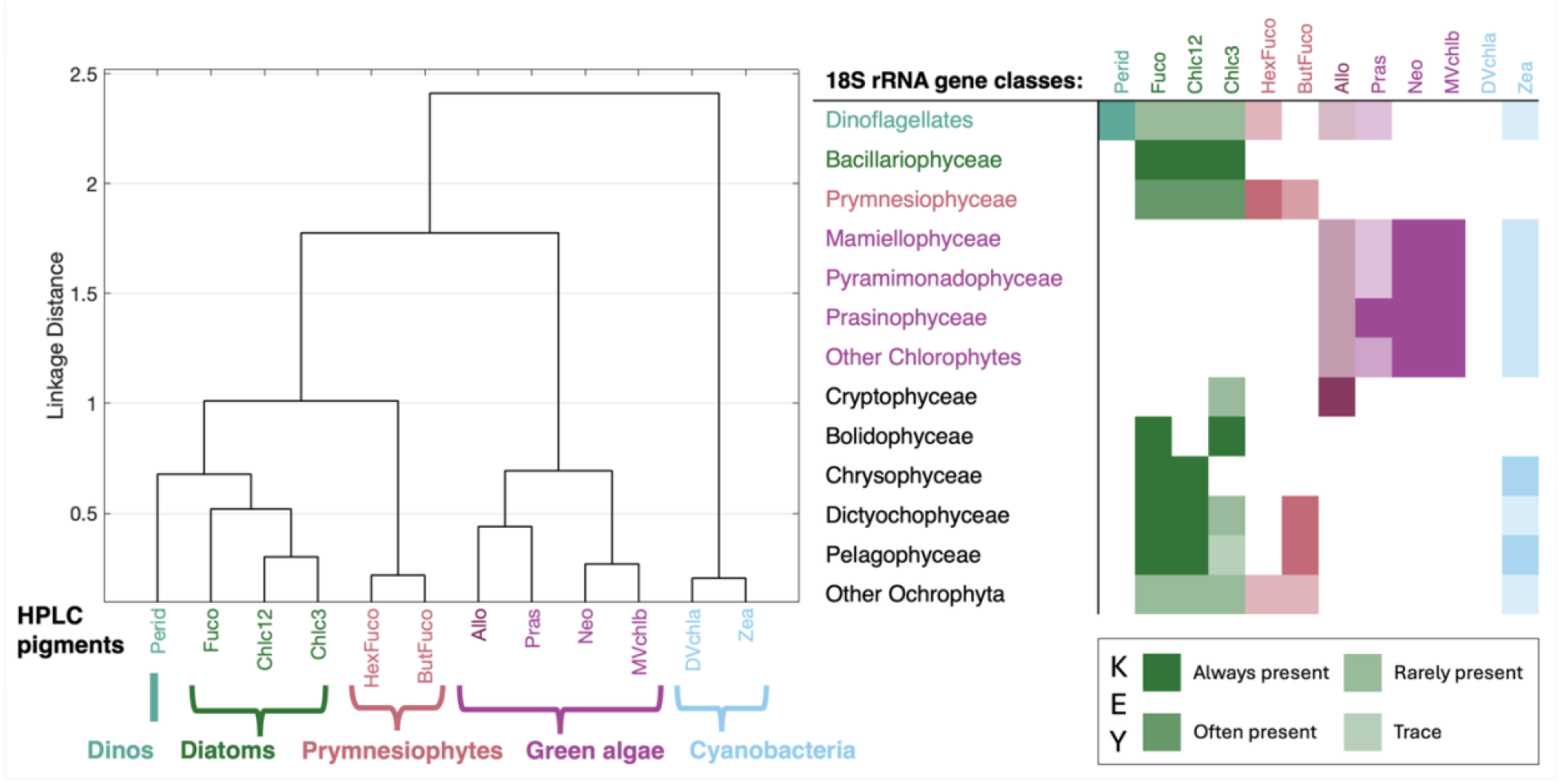
Phytoplankton groups considered in this study from both methods. Dendrogram showing the co-variability between thirteen phytoplankton pigments clustered into five phytoplankton groups (teal = dinoflagellates, green = diatoms, pink = prymnesiophytes, purple = green algae, light blue = cyanobacteria). Class-level groupings from the 18S rRNA gene data are shown, colored by the corresponding phytoplankton pigment cluster. Shading intensity indicates the likelihood of that pigment occurring in that class (e.g., Dictyochophytes always contain Fuco, Chlc12, and ButFuco, often contain Chlc3, and rarely contain Zea) (Jeffrey et al. 2011; Kramer and Siegel 2019).

### 2.4 18S rRNA gene sequences

Finally, phytoplankton taxonomy was also evaluated from 18S rRNA gene sequences. This gene was selected to describe the whole eukaryotic photosynthetic phytoplankton community, including dinoflagellates, which are a diverse phytoplankton group that cannot fully be quantified or described using 16S rRNA genes (Lin 2011). In order to maximize public data availability between *R_rs_*(*λ*)-modeled pigments, HPLC pigments, and sequence data, this study focused on 18S rRNA gene sequences instead of other genetic markers. The relative abundance of 18S rRNA gene sequences is broadly correlated with relative phytoplankton abundance measured across other methods for most broad taxonomic groups within the eukaryotes (Lin et al. 2019; Catlett et al. 2020, 2022; Kramer et al. 2024a) (see, for this study, e.g., **Figure S2**), but cyanobacteria are prokaryotes and are not targeted by 18S rRNA gene sequencing, so the sequence data used in this study do not describe prokaryotic phytoplankton. Since HPLC pigments were also measured from all of these samples, the average Zea/Tchla and DVchla/Tchla within each sample could be used to describe the relative contribution of cyanobacterial pigments, but this metric could not be validated with DNA data.

The specific methods for gene amplification and sequencing for all datasets used in this study have already been described in detail elsewhere and therefore are summarized in the Appendix. The number of photosynthetic phytoplankton ASVs detected in a given dataset varied by two orders of magnitude among datasets, and thus were summed into thirteen major shared taxonomic classes for consistency across datasets and for comparison of the relative contributions of phytoplankton diversity with the twelve accessory pigments (**Figure 2**): Dinophyceae (dinoflagellates); Mamiellophyceae, Pyramimonadophyceae, Prasinophyceae, other Chlorophytes (green algae); Prymnesiophyceae (prymnesiophytes); Bacillariophyceae (diatoms); Bolidophyceae; Chrysophyceae; Cryptophyceae; Dictyochophyceae; Pelagophyceae; and other Ochrophyta.

### 2.5 Phytoplankton carbon (C_phyto_)

Phytoplankton carbon (*C_p_*_ℎ*yto*_) was calculated from total chlorophyll-*a* concentration following (Graff et al. 2015) as a proxy for phytoplankton productivity. *C_p_*_ℎ*yto*_ was also calculated from a superior proxy, particulate backscattering at 470 nm (*b_bp_*(470)) (Graff et al. 2015), for NAAMES and EXPORTS, which were the two cruises for which these data were publicly available in the same times, places, and depths as HPLC pigment concentrations and 18S rRNA gene sequences in this global dataset.

### 2.6 Shannon diversity index

To assess phytoplankton diversity (as a factor of both richness and evenness), the Shannon-Weaver diversity index (Kim, Bo-Ra et al. 2017) was calculated for: (1) relative concentrations of 12 phytoplankton pigments to total chlorophyll-*a* (prokaryotic and eukaryotic phytoplankton), (2) relative sequence abundances of 13 18S rRNA gene classes (eukaryotic phytoplankton only; **Figure 1C**), (3) relative SDP-modeled phytoplankton pigment concentrations to modeled total chlorophyll-*a* concentration (prokaryotic and eukaryotic phytoplankton), and (4) relative sequence abundances for ASV-level 18S rRNA genes (eukaryotic phytoplankton only).

### 2.7 Network-based community detection analysis

To assess global-scale trends in phytoplankton community composition between these different methods, samples were divided into distinct communities using network-based community detection analysis following (Kramer et al. 2020, 2024b). Full details of this approach are provided in the Appendix. Following the separation of samples into communities, phytoplankton taxonomy in each pigment community or 18S rRNA gene community was also evaluated based on the mean phytoplankton composition within that community (**Figure S3**). Given the limitations of comparing taxonomy between HPLC pigments and the 18S rRNA gene, these communities were not assigned putative taxonomy and were instead described by number (1, 2, or 3).

### 2.8 Statistical analysis

In addition to the analysis described above, an ANOVA was performed to assess the statistical similarity between communities separated from all samples in the HPLC and 18S rRNA datasets (N = 324). Accessory pigment concentrations were normalized to Tchla and the similarity of pigment ratios between and among communities was assessed. The *P* value was calculated for each ANOVA and is reported in the text. All statistical analysis was performed in MATLAB 2024b.

## 3. Results

### 3.1 Patterns in global surface ocean phytoplankton community composition from pigments and 18S rRNA genes are similar

HPLC pigments and 18S rRNA gene sequences provide different taxonomic resolution for describing phytoplankton communities (**Figure 2**). While 12 distinct accessory pigments were used in this study, many of those pigments were shared within one taxonomic group (e.g., Fuco, Chlc12, and Chlc3 clustered in diatoms) and were not distinct markers, so only five distinct phytoplankton groups could be separated from HPLC pigments. For instance, the pigments Pras, Neo, and MVchlb are all found in green algae, and thus clustered together as a group of green algal accessory pigments that separated as a broad phytoplankton group with Allo, which is found in cryptophytes. From 18S rRNA gene sequences, however, three specific green algal classes were separated (Mamiellophyceae, Pyramimonadophyceae, Prasinophyceae). Other green algal classes were less common in this dataset and those ASVs were combined as “other Chlorophytes.” Cryptophytes separated distinctly in the 18S rRNA gene data from the four green algal classes. Other major phytoplankton classes can be better described by just one 18S rRNA gene class and one accessory pigment (e.g., dinoflagellates). Conversely, HPLC pigments provide a way to detect cyanobacteria based on the concentrations of Zea and DVchla (which is only found in *Prochlorococcus*), but the 18S rRNA gene is not found in prokaryotic phytoplankton. Despite the broad differences in taxonomic resolution between methods, there was generally good agreement between relative pigment concentrations (normalized to total chlorophyll-*a*) and relative 18S rRNA gene sequence abundances across groups for which these methods could be directly compared (**Figure S2** demonstrates a range of positive, significant linear correlations between methods, with the strongest relationship found for diatoms), as expected from prior studies (Kramer et al. 2024a).

Given the differences in the taxonomic resolution of phytoplankton composition detected by each method, it seemed likely that different communities would be retrieved between the two approaches. However, similar communities were separated by both HPLC pigments and 18S rRNA gene sequences (**Figure 3**), with similarities in both the dominant phytoplankton pigment composition and the sample distribution detected across the two datasets. Three communities were retrieved from the HPLC pigment data (**Figure 3A**). There was strong separation between communities, with a network modularity value of 0.33. The average concentration of Fuco/Tchla in HPLC Community 1 was significantly higher than in the other two communities; similar results were found for both Zea/Tchla and DVchla/Tchla in HPLC Community 2 and MVchlb/Tchla in HPLC Community 3 (ANOVA; *P* value <0.001).

**Figure 3.**
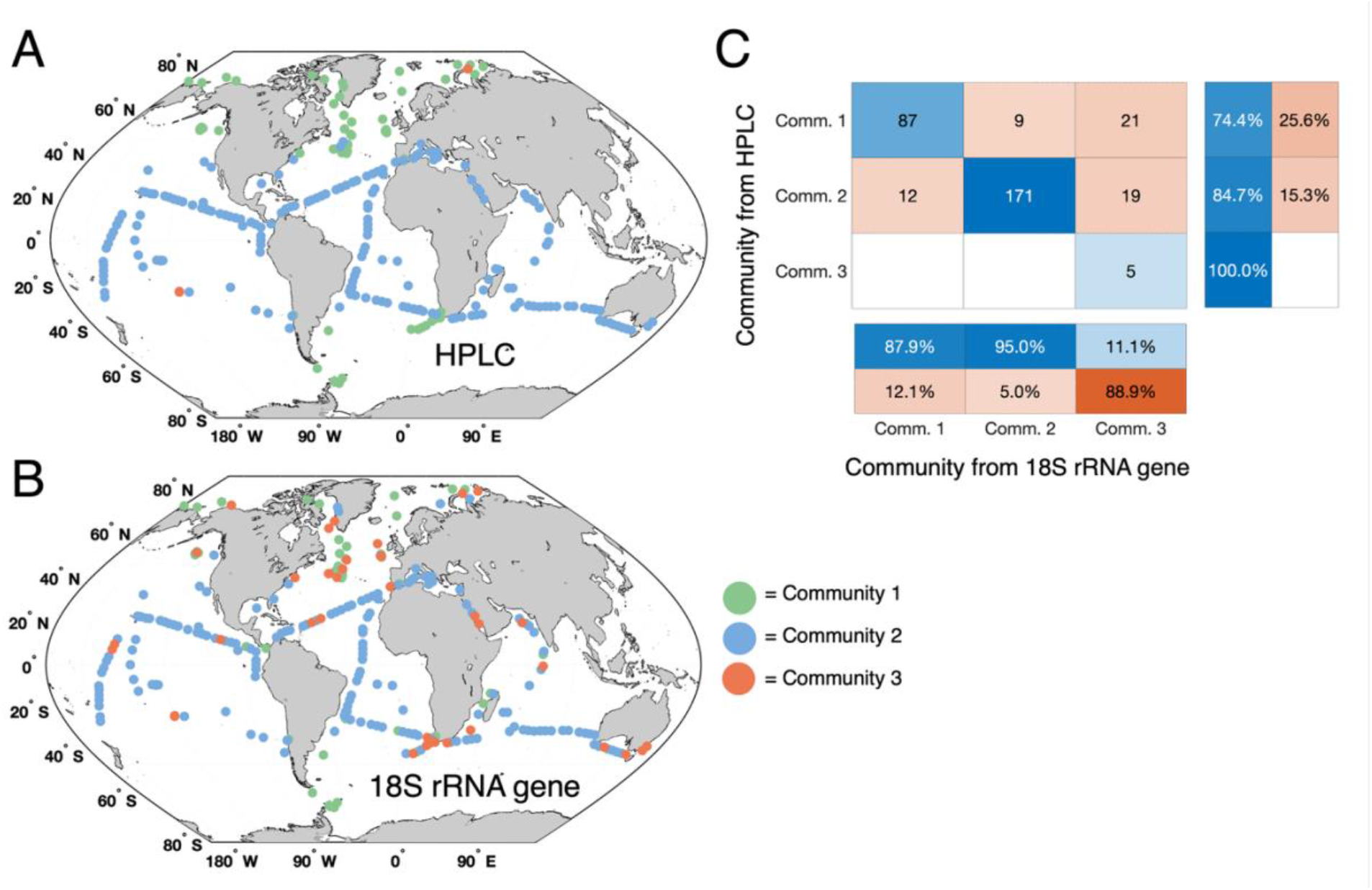
Global distribution of network-based community detection analyses for (A) relative HPLC pigment concentrations and (B) 18S rRNA gene relative class abundances (green = Community 1, light blue = Community 2, orange = Community 3). A confusion matrix (C) shows the comparison between the HPLC pigment-based communities and 18S rRNA gene-based communities. The number of “correctly” (shades of blue) vs. “incorrectly” (shades of red) assigned samples are given by numbered boxes, assuming that pigment-based communities are “correct.” The percentage of samples that are correctly (blue) vs. incorrectly (red) assigned for each method are also shown.

There were also three communities retrieved from the distinct 18S rRNA gene community detection analysis (**Figure 3B**), but with slightly less robust separation between communities (modularity value = 0.28). Since the 18S rRNA gene cannot be used to assess cyanobacteria, but HPLC pigments were also measured from these samples, a comparison of the average accessory pigment composition determined that these three 18S rRNA gene communities had similar mean pigment composition to the three HPLC communities (**Figure S3**). Specifically, these comparisons were performed by, e.g., assessing the mean pigment composition in all samples assigned to HPLC Community 1 (which includes both HPLC and 18S rRNA gene sequence data for each sample) and comparing that to the mean pigment composition in all samples assigned to 18S rRNA gene Community 1 (which also includes both HPLC and 18S rRNA gene sequence data for each sample). The mean Fuco/Tchla concentration and the mean relative sequence abundance of Bacillariophyceae were significantly different for the 18S Community 1, while the mean MVchlb/Tchla concentration and mean relative sequence abundance of all summed Chlorophyte classes were significantly different for 18S Community 3 (ANOVA; *P* value <0.001). The mean Zea/Tchla and DVchla/Tchla ratios for 18S Community 2 were also significantly different from the other 18S communities (ANOVA; *P* value <0.001).

Overall, when the HPLC communities and 18S rRNA gene communities were compared directly, 81% of all samples (262 of 324 total) were assigned to the same community (**Figure 3C**). The geographic distributions of communities defined by these two methods had broad similarities across the globe with some notable deviations (**Figure 3A-B**). The equatorial regions and subtropical gyres were dominated by Community 2 from both analyses, while the coasts and high latitude regions contained mostly Community 1. Only 5 samples were separated into Community 3 from the HPLC pigment data, while 45 samples were separated into Community 3 from 18S rRNA gene sequences. This direct comparison had the most disagreement between methods, with nearly 90% of the 18S Community 3 samples assigned to either HPLC Community 1 or 2 (**Figure 3C**). Since the MVchlb/Tchla ratio was significantly high in Community 3, the mean MVchlb/Tchla ratio for the 40 samples that disagreed was assessed. For those samples, this ratio was significantly lower than the mean MVchlb/Tchla ratio in HPLC Community 3 (0.10 vs. 0.30) but significantly higher than the mean MVchlb/Tchla ratio in all other samples (0.10 vs. 0.06; ANOVA, *P* value < 0.001). Taken together, the global trends in phytoplankton communities separated by HPLC pigments vs. 18S rRNA gene sequences were the same for the vast majority of samples and rare disagreements between methods highlighted opportunities for further analysis.

### 3.2 The shape of the relationship between ocean phytoplankton diversity and carbon biomass is unimodal across methods

By comparing across similar proxies for the different methods, the relationship between ocean phytoplankton diversity and carbon biomass was investigated (**Figure 4**). This relationship can be considered similar to the PDR, though here a stock was examined (carbon biomass) instead of a rate (productivity). First, this relationship was investigated with optically-derived data to consider the shape of the relationship if observations were limited to those we could make from space. Specifically, for this analysis, the Shannon diversity index was calculated from 12 phytoplankton pigment ratios modeled from hyperspectral *R_rs_*(*λ*) vs. phytoplankton carbon derived from *R_rs_*(*λ*)-modeled Tchla (**Figure 4A**). The shape of the this relationship closely matched the expected, unimodal distribution previously compiled for resampled phytoplankton microscopy data (Vallina et al. 2014). The shape was described by overlaying a generic Gaussian function, to capture the trend between variables. At low-to-medium carbon biomass concentrations, phytoplankton pigment diversity increased across a gradient in carbon biomass, before reaching a plateau. Since not all samples in the HPLC + *R_rs_*(*λ*) dataset (Section 2.2) were included in the global HPLC dataset used here (Section 2.3), the samples shared between both datasets were simply colored by their HPLC community identity. Samples in HPLC Community 2 were found most commonly at lower carbon biomass concentrations in the relationship between *R_rs_*(*λ*)-modeled diversity and carbon biomass, while samples in HPLC Community 1 were mostly found at higher carbon biomass concentrations.

**Figure 4.**
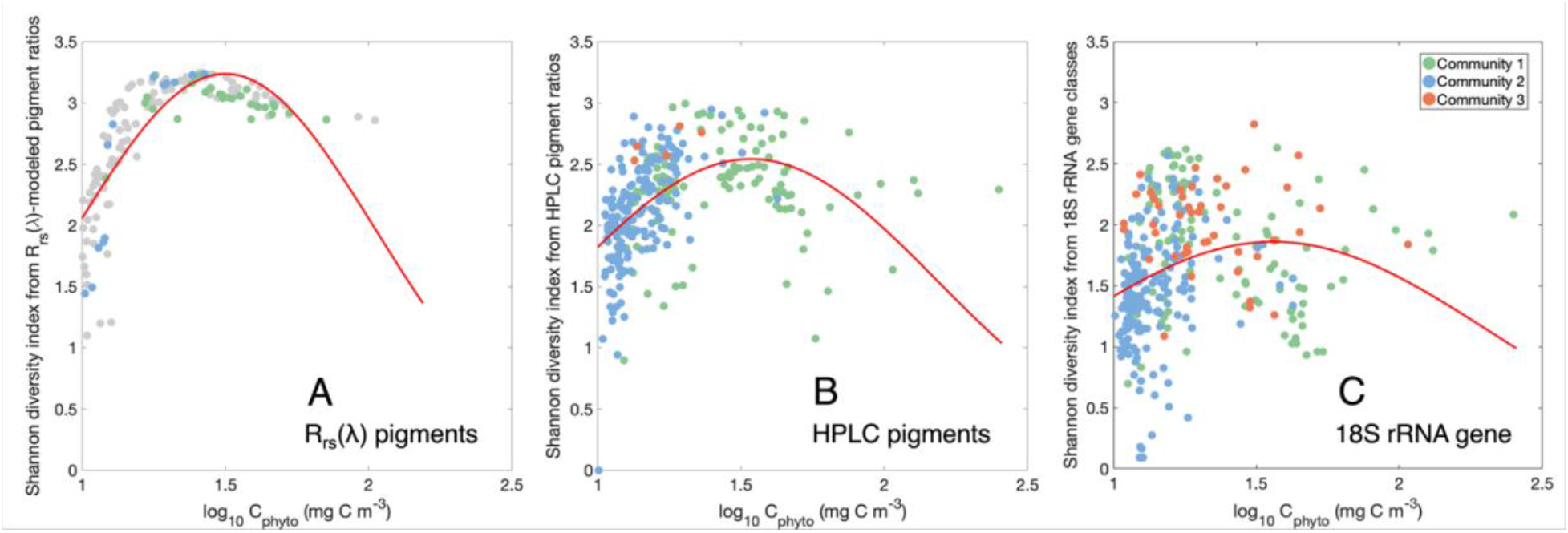
Relationship between phytoplankton diversity based on the Shannon index for (A) ratios of remote sensing reflectance (*R_rs_*(*λ*))-derived pigments to *R_rs_*(*λ*)-derived total chlorophyll-*a* (N = 162), (B) ratios of HPLC pigments to total chlorophyll-*a* (N = 324), and (C) relative abundances of 18S rRNA gene classes (eukaryotic phytoplankton) vs. phytoplankton carbon biomass from total chlorophyll-*a* for all panels (N = 324), following (Graff et al. 2015). All samples are colored by the community defined by network-based community detection analysis for each method and shown in Figure 3A (HPLC) and 3B (18S rRNA genes). For both methods, green = Community 1, light blue = Community 2, orange = Community 3. Only 41 samples in the HPLC-*R_rs_*(*λ*) dataset are shared with the HPLC-18S rRNA gene dataset, so those are the only samples colored by community in panel (A).

While assessing the relationship between phytoplankton diversity and carbon biomass from remotely-sensed variables will ideally encourage future evaluations with satellite-derived variables, the relationship was also assessed with the larger, paired in situ dataset to compare trends. The shape of this relationship with diversity calculated from both HPLC pigments (**Figure 4B**) and 18S rRNA gene sequences (**Figure 4C**) matched the unimodal distribution found in **Figure 4A**. Generic Gaussian functions were also fit to the data, to capture the broad trend between variables (**Table 1**). A normal distribution was the best fit for all variables except the ASV-level 18S rRNA gene diversity (Figure 5B), which was fit with a split Gaussian (amplitude = 5.98, center = 0.99, left width = 12.2, right width = 1.33; *R^2^* = 0.06).

**Figure 5.**
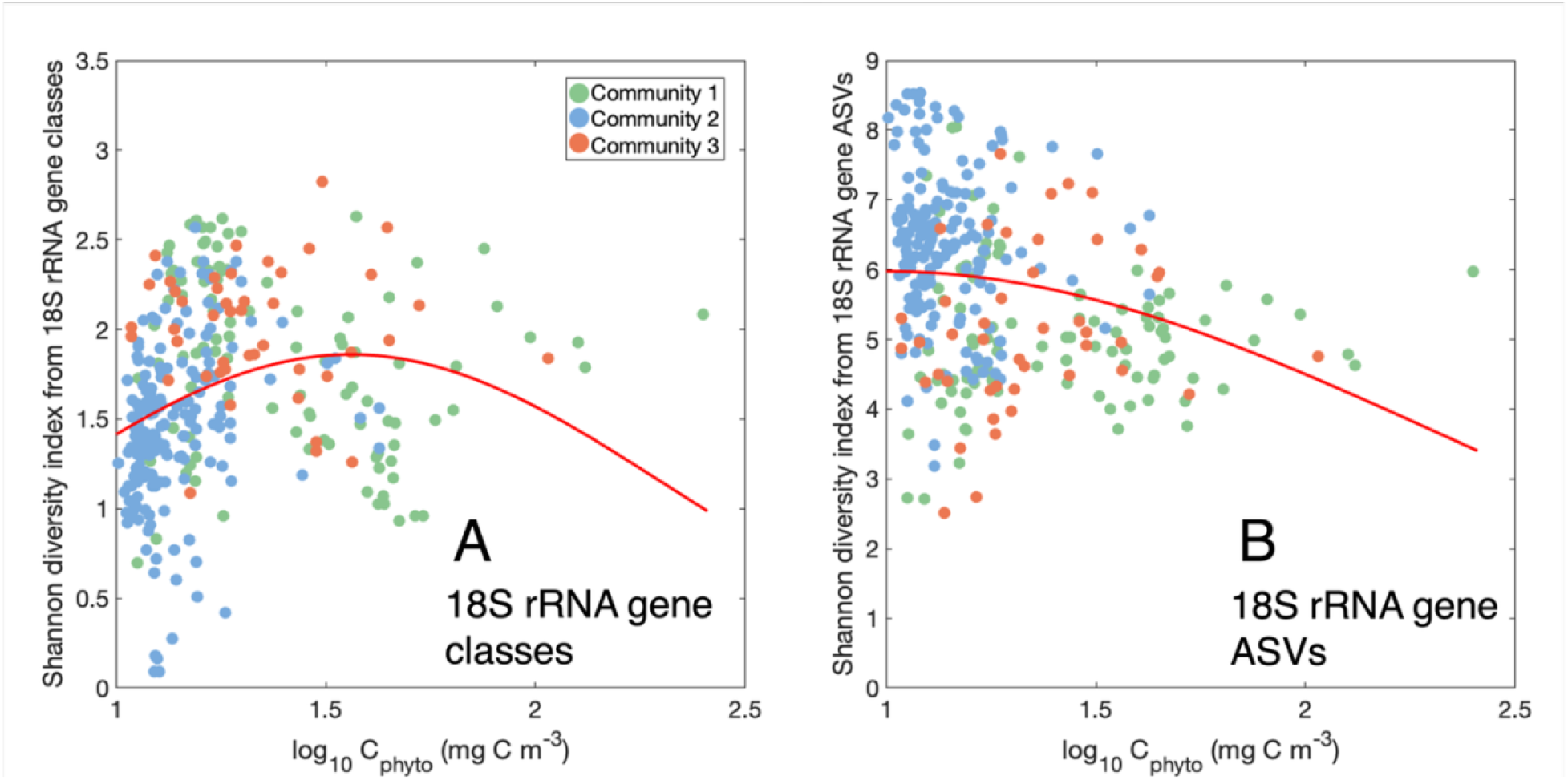
Relationship between eukaryotic phytoplankton diversity based on the Shannon index for (A) class-level relative abundances of 18S rRNA gene data (N = 324), and (B) ASV-level relative abundances of 18S rRNA gene data vs. phytoplankton carbon biomass (N = 324; from total chlorophyll-*a* for all panels, following (Graff et al. 2015)). All samples are colored by the community defined by the 18S rRNA gene network-based community detection analysis and shown in Figure 3B (green = Community 1, light blue = Community 2, orange = Community 3).

**Table 1.**
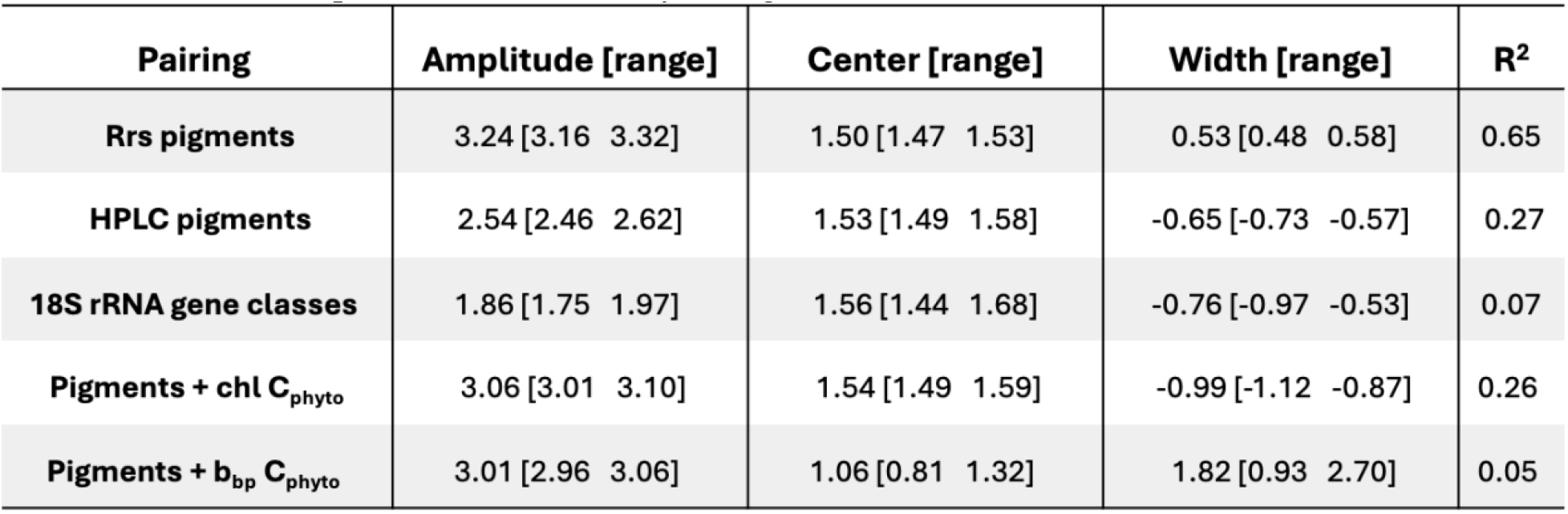
Fitted model parameters, uncertainty, and goodness of fit for all normal Gaussian functions.

| **Pairing** | **Amplitude [range]** | **Center [range]** | **Width [range]** | **R<sup>2</sup>** |
| --- | --- | --- | --- | --- |
| **Rrs pigments** | 3.24 [3.16 3.32] | 1.50 [1.47 1.53] | 0.53 [0.48 0.58] | 0.65 |
| **HPLC pigments** | 2.54 [2.46 2.62] | 1.53 [1.49 1.58] | -0.65 [-0.73 -0.57] | 0.27 |
| **18S rRNA gene classes** | 1.86 [1.75 1.97] | 1.56 [1.44 1.68] | -0.76 [-0.97 -0.53] | 0.07 |
| **Pigments + chl C<sub>phyto</sub>** | 3.06 [3.01 3.10] | 1.54 [1.49 1.59] | -0.99 [-1.12 -0.87] | 0.26 |
| **Pigments + b<sub>bp</sub> C<sub>phyto</sub>** | 3.01 [2.96 3.06] | 1.06 [0.81 1.32] | 1.82 [0.93 2.70] | 0.05 |

The highest diversity was found at intermediate carbon biomass concentration, with relatively lower diversity found at both the lowest and highest carbon biomass concentrations. There was also a trend with the broad phytoplankton community determined by network-based community detection analysis that matched the pattern found in **Figure 4A**: samples in HPLC Community 2 were concentrated at lower carbon biomass concentrations and samples in HPLC Community 1 were mostly found at higher carbon biomass concentrations, though this latter community was more evenly distributed across a gradient in carbon biomass concentration (**Figure 4B**). The peak in diversity was a hinge point between these two communities, with HPLC Communities 2 and 1 shifting at intermediate carbon biomass concentration.

Meanwhile, a similar trend existed for the 18S rRNA gene communities, with 18S Community 2 mostly found at lower carbon biomass concentrations and 18S Community 1 mostly found at higher concentrations (**Figure 4C**). However, 18S Community 2 was found more at higher carbon biomass concentrations at times in this dataset, and 18S Community 3 (which contained more samples than HPLC Community 3; **Figure 3C**) was distributed across a range of concentrations. At the highest diversity levels, a range of conditions was found, all representing different phytoplankton communities with intermediate carbon biomass. On a regional scale, the shape of this relationship often captured either one side of the unimodal curve or the other, rather than representing the whole spectrum (**Figure S4**). For instance, from the tropical/subtropical Malaspina dataset (**Figure S1**), the relationship between diversity and carbon biomass was monotonically increasing, while the relationship across the subarctic EXPORTS dataset showed the opposite trend, particularly from 18S rRNA gene sequences (**Figure S3B**).

### 3.3 Proxies are important: different metrics for diversity and carbon biomass change the shape of the relationship between phytoplankton diversity and carbon biomass

The similar patterns between phytoplankton diversity derived from pigments (prokaryotes and eukaryotes; **Figure 4A-B**) and the 18S rRNA gene (eukaryotes only; **Figure 4C**) vs. carbon biomass suggest correspondence across methods when phytoplankton diversity was compared at a similar level of taxonomic complexity (e.g., 12 accessory pigments to Tchla vs. 13 18S rRNA gene classes). This taxonomic resolution was chosen to match the number of variables available from satellite-based estimates of phytoplankton diversity (12 *R_rs_*(*λ*)-modeled accessory pigment concentrations; **Figure 4A**).

However, gene sequence data has much higher taxonomic resolution than the 13 photosynthetic phytoplankton classes shown here. The Shannon index was calculated again for each sample using ASV-level 18S rRNA gene data, which contained hundreds to thousands of ASVs depending on the sample (**Figure 5**). Much of the broad shape of the relationship observed between diversity and carbon biomass at the class-level (**Figure 4C**) was obscured when diversity was defined by ASV-level 18S rRNA gene data (**Figure 5B**). The Shannon index was also much higher from ASV-level data (values ranged from 3-9) than class-level data (values from 0-3), as would be expected given the increase in alpha diversity. 18S Community 2 was also distributed differently when diversity was calculated at the ASV-level instead of the class-level, with many more examples of high diversity at low carbon biomass concentrations, matching past observations from global surveys using high resolution genomic sequencing (Ibarbalz et al. 2019). These samples also likely contain high relative contributions of cyanobacteria that could not be observed with the 18S rRNA gene sequences (**Figure S3**), raising further questions about invisible diversity between methods, and whether that taxonomic diversity corresponds to functional and/or genetic diversity (Biller et al. 2015).

Similar to the importance of the diversity metric, the metric used for phytoplankton carbon biomass may also influence this relationship. The similar patterns found here (**Figure 4**) used the same metric for phytoplankton carbon (*C_p_*_ℎ*yto*_), which was derived from total chlorophyll-*a* concentration here. Although this proxy was chosen to maximize the number of paired samples in this global dataset, it has been previously demonstrated (Graff et al. 2015) that the relationship between in situ *C_p_*_ℎ*yto*_ and total chlorophyll-*a* is not as robust as the relationship between in situ *C_p_*_ℎ*yto*_ and particle backscattering at 470 nm (*b_bp_*(470)). Thus, for a subset of samples in this dataset for which HPLC pigment concentrations and *b_bp_*(470) were measured concurrently (N =), the shape of the PDR was compared with consistent diversity information (Shannon index from 12 ratios of accessory pigments to Tchla) but different proxies for carbon biomass (**Figure 6**). Additional surface samples with these two data types from NAAMES and EXPORTS were also included relative to the global dataset used otherwise in this analysis (Kramer et al. 2020; Siegel et al. 2021; Johnson et al. 2023). Some similar groups of samples with low diversity and low biomass or high diversity and low biomass emerged between methods (**Figure 6**). Similarly, the peak in diversity coincided with intermediate phytoplankton carbon biomass values for both approaches.

**Figure 6.**
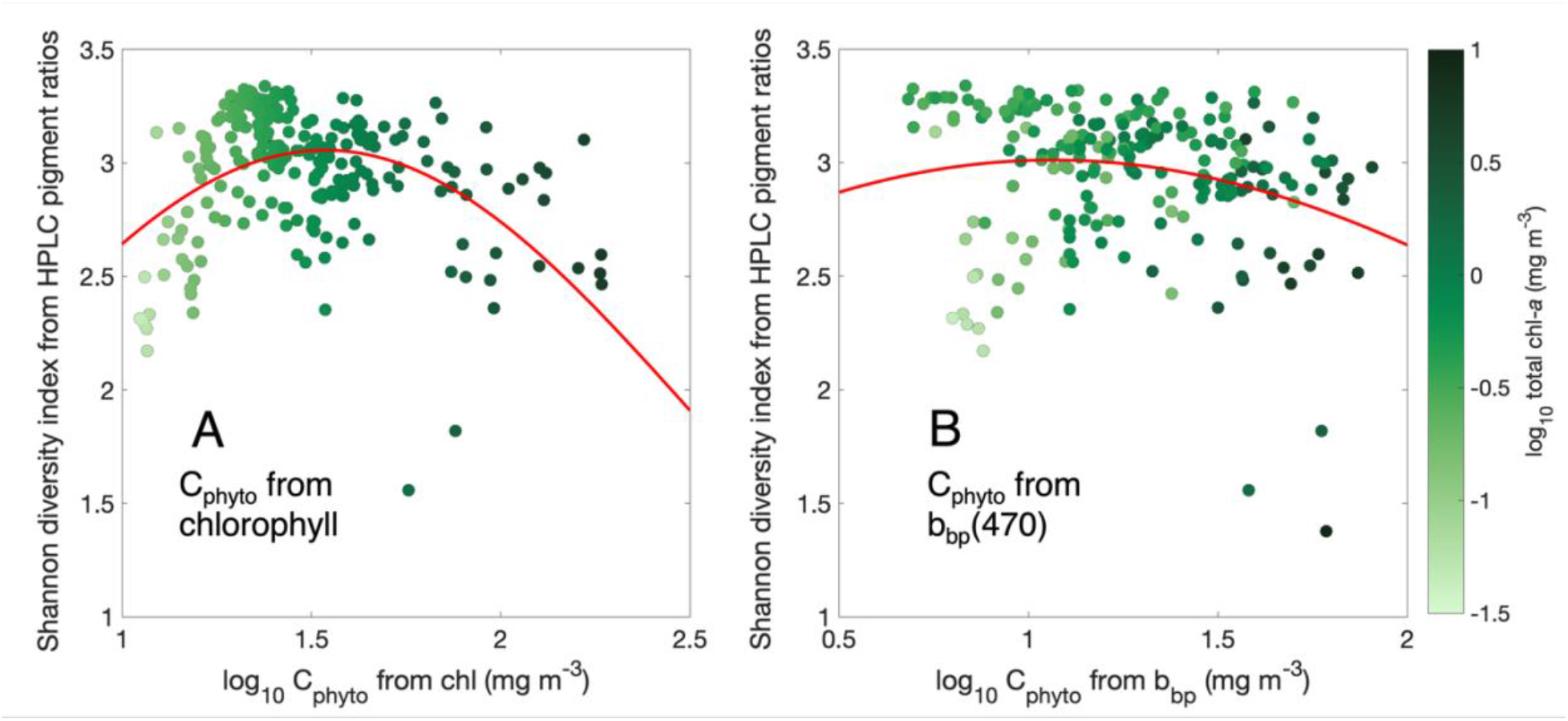
Relationship between prokaryotic and eukaryotic phytoplankton diversity based on the Shannon index for HPLC pigments and phytoplankton carbon biomass on NAAMES and EXPORTS from (A) total chlorophyll-*a* and (B) particle backscattering (*b_bp_*(470), following (Graff et al. 2015)). All samples are colored by total chlorophyll-*a* concentration from HPLC.

However, there were several samples with high diversity and low phytoplankton biomass from the *b_bp_*(470)-based proxy (**Figure 6B**), because *C_p_*_ℎ*yto*_ was overestimated from chlorophyll-*a* for those samples (**Figure 6A**). The conflation of chlorophyll-*a* concentration and phytoplankton biomass has been well documented for both in situ sampling and remote sensing applications (Behrenfeld and Boss 2006). Chlorophyll-*a* concentration is more often measured in conjunction with biological samples (e.g., 18S rRNA gene sequences) than *b_bp_*(470), allowing easier comparisons between datasets but perhaps less accurate proxies in the open ocean for broader Earth system variables such as phytoplankton carbon.

## 4. Discussion

Characterizing the relationship between phytoplankton diversity and carbon biomass is essential for a complete understanding of Earth system processes, from biogeochemical cycling to marine food webs to carbon sequestration and climate. Many models that target a mechanistic understanding of these processes will incorporate a phytoplankton diversity parameter, often limited to a small number of phytoplankton functional types or size classes. These models aim to simulate the impact of multiple phytoplankton taxa on an ecosystem to understand the role of phytoplankton in mediating climate, or the impact of a changing climate on phytoplankton and other related components of the Earth system, such as carbon sequestration and nutrient cycling (Le Quéré et al. 2005; Gregg and Casey 2007; Dutkiewicz et al. 2009). The results shown here demonstrate that the number and types of phytoplankton groups used to describe diversity affected the resulting relationships between phytoplankton diversity and carbon biomass, which is related to ecosystem productivity. Similarly, the shape of this relationship changed based on the calculation used for phytoplankton carbon biomass, as chlorophyll-*a* concentration is an imperfect proxy. Ultimately, both the resolution of the phytoplankton taxonomic data used to calculate a diversity index and the proxy used to derive phytoplankton carbon had an effect on the shape and magnitude of the relationship between these variables in the global surface ocean.

While this study evaluated global-scale variations in a stock (phytoplankton carbon biomass) rather than a rate (ecosystem productivity), these metrics have long been used interchangeably to assess the shape of the PDR (Irigoien et al. 2004; Vallina et al. 2014; Ibarbalz et al. 2019; Lampe et al. 2025). Thus, if the relationship presented here is comparable to the PDR, the significance of the associations between phytoplankton diversity and ecosystem productivity can be considered. When evaluating the role of the PDR in structuring essential Earth system processes, the scale of inquiry becomes particularly relevant. This question of scale can be assessed in two ways, both of which were examined in this study: first, as the global vs. regional geographic scale on which the analysis was conducted, and second, as the taxonomic resolution at which the PDR was constructed. Ultimately, both of these considerations were relevant for defining the shape of the PDR, and both are important for selecting the best method to assess the global ocean and construct the PDR. The in situ HPLC pigment and 18S rRNA gene sequence data used here provided global coverage when compiled across major field campaigns that took over a decade to complete, with samples collected from 2009-2021 across datasets. When only one field campaign was considered at a time, the shape of the relationship captured one side or the other of the classic unimodal curve found in a global dataset (Acevedo-Trejos et al. 2018; Ibarbalz et al. 2019). For instance, the Malaspina expedition sampled a limited latitude range that corresponded to a limited range of carbon biomass concentrations and associated phytoplankton diversity, leading to a positive slope; alternately, the EXPORTS field campaigns sampled at similar subpolar latitudes but in two different basins and ecosystem states, leading to a negative slope. This scale-dependence has also been demonstrated from observations and models of the PDR on land (Chase and Leibold 2002; Gonzalez et al. 2020).

Additionally, the scale of the phytoplankton taxonomic resolution is highly relevant for defining the shape of this relationship. The relationship between phytoplankton diversity and carbon biomass was roughly similar when assessed from phytoplankton pigments and class-level 18S rRNA gene sequence data, despite the slightly different communities assessed by each method (prokaryotes + eukaryotes vs. eukaryotes only) but changed for the much higher resolution ASV-level 18S rRNA genes. The assessment of taxonomy from the 18S rRNA gene can also be affected by the hypervariable region that was sequenced, with similar but slightly different taxonomy retrieved between, e.g., V4 vs. V9 (Stoeck et al. 2010; Zimmermann et al. 2024). Finally, none of the methods used in the present study were able to assess cell size, which is also an important consideration for both carbon biomass stocks and productivity rates. While the taxonomic resolution varied between the 5 phytoplankton groups separated by HPLC pigments and the 13 classes separated by 18S rRNA genes, three similar communities were retrieved from these two approaches across the global dataset, with the same community assignment for over 80% of the samples. These similar trends between methods are promising for describing much of the global ocean with a consistent metric for phytoplankton community composition.

If one goal of an Earth system model is to accurately capture as much of the globe as possible, then the results presented here demonstrate that the method selected to assess the shape of the PDR for model applications should maximize both geographic scope (global coverage) and phytoplankton taxonomic resolution. For global monitoring of the surface ocean, no in situ sampling can match the spatiotemporal resolution of remote sensing approaches, while no satellite measurement can come close to the taxonomic resolution of DNA metabarcoding. Ocean color satellites provide a global composite approximately every 8 days and survey the optical features of phytoplankton communities to provide relevant proxies for both taxonomy (via modeled phytoplankton pigments, available only from hyperspectral *R_rs_*(*λ*) as measured globally by PACE; (Kramer et al. 2022; Cetinić et al. 2024)), carbon biomass (Graff et al. 2015; Fox et al. 2022), and productivity (Behrenfeld and Falkowski 1997). Although 18S rRNA gene sequence data (and other molecular approaches) can describe the phytoplankton community with more taxonomic depth, methods do not *currently* exist to convert between phytoplankton taxonomy at this resolution and the coarser pigment-based groups modeled from satellite remote sensing data collected at high spatial resolution.

This work demonstrates that combining global-scale genomic sampling with pigment-level phytoplankton taxonomic resolution can still represent similar relationships between phytoplankton community composition and diversity vs. carbon biomass, providing the potential to explore these relationships over time in the PACE data record at the scales needed to evaluate global change. However, ocean color satellites are only measuring the upper layer of the sunlit ocean, and significant phytoplankton carbon biomass and productivity occur at depth and across stratified water masses.

Satellite-based approaches can only capture the diversity of upper ocean phytoplankton when mixed layer depths are deeper than euphotic depths, and recent work has shown that the euphotic depth is often much deeper than previously assumed, particularly when defined as the region of the ocean in which photosynthesis is possible (Wu et al. 2021). Given the strengths and limitations of the various methods assessed here, the strongest future evaluations of these global relationships will come from combining methods that consider broad spatiotemporal scales (i.e., ocean color remote sensing) as well as high taxonomic resolution (i.e., DNA metabarcoding). Methods that bridge these two approaches and link the phytoplankton community to ocean color (i.e., HPLC phytoplankton pigments) will remain important for establishing strong relationships across methods in space and time, across regions and seasons.

The high degree of variability found between and among methods for assessing phytoplankton diversity and carbon biomass in this study also highlights the importance of global monitoring using consistent approaches in a changing world. As the ocean changes in response to anthropogenic climate change, phytoplankton diversity and productivity will naturally be affected (Behrenfeld 2014; Behrenfeld et al. 2015). The palaeoceanographic record suggests that the impact of climate on phytoplankton (and vice versa) can be extreme (Falkowski and Oliver 2007). The strong latitudinal structuring of the PDR in terrestrial systems (Pärtel et al. 2007) and the correlations between diversity and temperature in marine systems (Ibarbalz et al. 2019; Anderson et al. 2021) suggest that climate change will also have strong impacts on the relationships between phytoplankton diversity, carbon biomass, and productivity across regions and ecosystems, in response to changing temperature and ocean acidity (Dutkiewicz et al. 2015; Bestion et al. 2021). Regional modeling approaches also suggest that the diversity of phytoplankton may increase in certain ecosystems under future warming scenarios, and that the increase could be more pronounced than predicated changes in chlorophyll-*a* concentration, again demonstrating the importance of scale (Dutkiewicz et al. 2019; Henson et al. 2021). These results cumulatively indicate that the response of a given phytoplankton community to a changing ocean is highly variable. Surveying changes in diversity, carbon biomass, *and* productivity, as is newly possible with PACE, will allow for an understanding of the phytoplankton community underlying the global distribution of chlorophyll-*a*. Evaluating the shape of these variable relationships across methods and scales can only help to improve estimates of the future relationships in a warming world.

Assessing phytoplankton diversity and carbon biomass on global scales is also relevant for evaluating large-scale ecosystem function, both in situ and for modeling considerations. A diverse assemblage of phytoplankton with varied niches for temperature and nutrient regimes is expected to be more productive over a seasonal cycle and in a changing world: as temperature and acidity rise and/or as nutrient concentrations decrease, ecosystems with a diverse assortment of phytoplankton will remain more productive. Broadly, the range and richness of most phytoplankton taxa are threatened by climate change, due to changing physical and biogeochemical conditions which will limit the regions in which many taxa can grow and thrive (Ibarbalz et al. 2019; Benedetti et al. 2021; Chaffron et al. 2021).

Phytoplankton communities may become less diverse under future warming scenarios and ecosystems may lose essential functions as a result—such as the ability to efficiently sequester carbon via the biological pump. The strongest impacts on phytoplankton diversity are expected at high latitudes (Ibarbalz et al. 2019; Benedetti et al. 2021), where export is currently highest via the biological carbon pump (Boyd et al. 2019). Given the importance of specific phytoplankton taxa for predicting the magnitude of the biological pump (Armstrong et al. 2001; Guidi et al. 2016; Duret et al. 2020; Preston et al. 2020; Kramer et al. 2025), changes in phytoplankton diversity and ecosystem productivity will impact climate and carbon sequestration on broad scales. The community detection analysis performed here found consistent communities of phytoplankton across latitudinal gradients between two different methods, which were then similarly distributed in their relationship with phytoplankton carbon biomass, with distinct taxa and pigments found at different carbon biomass concentrations. However, assessing the phytoplankton community at varying taxonomic resolution resulted in a different relationship between these variables, so the magnitude of these shifts will be driven by the severity of potential diversity loss under climate change. If entire functional groups are lost as a result of rising ocean temperatures, these relationships might shift more drastically than with only limited ASV-level diversity loss. Regardless, as phytoplankton taxonomy, carbon biomass, and productivity change with a changing climate, so the shape of the PDR will change across the globe.

## Conclusions

This study demonstrated that data are needed from consistent methods across global scales to capture the relevant variability in phytoplankton diversity and carbon biomass. The approach of combining in situ and optical methods was motivated by the importance of understanding the relationships between phytoplankton diversity, carbon biomass, and productivity on broad spatial scales but with adequate taxonomic resolution. The datasets used here to assess this relationship across spatial scales represent over a decade of studies dedicated to collecting high-quality, paired samples across space and time in the global ocean. Unlike plants on land, phytoplankton in the ocean exist in a dynamic physical system that is mostly at steady state on long timescales. However, the system might not be at steady state in the moment that it is captured for a discrete sample. Assessing the true variability in phytoplankton biogeography on relevant timescales for phytoplankton requires data to be collected daily and globally, which is made possible by satellite remote sensing—though taxonomic resolution is lost at those spatiotemporal scales. By combining high-resolution metrics for phytoplankton community composition with remotely-sensible variables, this study aimed to consider the strengths and limitations of assessing phytoplankton diversity and carbon biomass from space. However, the current study was limited to the sunlit region of the ocean that is visible to ocean color satellites. Given that most carbon biomass in many regions of the ocean is found at depths below where a satellite can see, more work will be needed to evaluate the variability in the relationships shown here with depth resolution. Additionally, future studies could integrate information about zooplankton grazing and physical mixing (turbulence, etc.) to consider top-down vs. bottom up controls on the distribution of organisms across this unimodal relationship. Ultimately, the broad similarity in the relationship between phytoplankton diversity and carbon biomass across in situ and remotely-sensible approaches motivates future applications of these methods for global application, though caution is warranted in understanding how the level of taxonomic complexity, the proxy for carbon biomass, the relationship between biomass and productivity, and the sampling depth range will affect the resulting relationship between these variables.

## Conflict of Interest Statement

The author declares no competing interests.

## Author Contributions

SJK designed research, procured funding, performed analysis, created figures, and wrote the paper.

## Data Availability Statement

All data used in this study are publicly available. EXPORTS HPLC pigments and 18S rRNA gene sequences: https://seabass.gsfc.nasa.gov/experiment/EXPORTS. NAAMES HPLC pigments and 18S rRNA gene sequences: https://seabass.gsfc.nasa.gov/experiment/NAAMES. Tara Oceans and Tara Polar HPLC pigments: https://seabass.gsfc.nasa.gov/experiment/Tara_Oceans_expedition, https://seabass.gsfc.nasa.gov/experiment/Tara_Oceans_Polar_Circle. Tara 18S rRNA gene sequences: https://doi.org/10.5281/zenodo.7551643. Malaspina 18S rRNA gene sequences: https://doi.org/10.5281/zenodo.8363876 and HPLC pigments: https://doi.org/10.1594/PANGAEA.948819. GEOTRACES GP10 18S rRNA gene sequences: https://github.com/jcmcnch/Global-rRNA-Univeral-Metabarcoding-of-Plankton and HPLC pigments: https://doi.org/10.5285/ff46f034-f47c-05f9-e053-6c86abc0dc7e. SDP model code is available at: https://github.com/sashajane19/Rrs_pigments, https://github.com/max-danenhower/rrs-SDP-pigments, and https://nasa.github.io/oceandata-notebooks/notebooks/oci/oci_sdp.html.

## Acknowledgments

SJK was supported for much of this work by a Simons Foundation Postdoctoral Fellowship in Marine Microbial Ecology (LS-FMME-00986836) and by the David and Lucile Packard Foundation. Support was also provided by NASA Ocean Biology and Biogeochemistry (Grant #80NSSC25K7428). Thank you to everyone involved in funding, collecting, and distributing data from the field campaigns shown here— this publicly-available, high-quality oceanographic data is extremely valuable and very appreciated. SJK is grateful for conversations with Federico Ibarbalz, Emmanuel Boss, Lee Karp-Boss, and Colleen Durkin, who all provided helpful suggestions and comments that greatly improved this work. Two anonymous reviewers provided thoughtful, rigorous suggestions that were very helpful in revising this work.

## Appendix

### Supplemental Methods

#### 18S rRNA gene sequence methods

The V4 region of the 18S rRNA gene was sequenced and taxonomy was assigned to raw sequences using the same DADA2 workflow and the PR2 database for: the Tara Oceans and Tara Polar expeditions (Guillou et al. 2013; de Vargas et al. 2015; Callahan et al. 2016; Delage et al. 2023), Malaspina (Junger et al. 2023), EXPORTS (SeaBASS 2018; Kramer et al. 2025), and GEOTRACES GA10 (McNichol et al. 2025a). In the NAAMES dataset, the V9 region of the 18S rRNA gene was sequenced and taxonomy was assigned using the PR2 database (SeaBASS 2014; Kramer et al. 2024a). Phytoplankton taxonomy has been found to be comparable at the class level between 18S rRNA V4 and 18S rRNA V9, so these samples were included in the global dataset (Zimmermann et al. 2024). After taxonomy was assigned, individual DNA sequences (amplicon sequence variants; ASVs) were assigned as either photosynthetic or heterotrophic taxa in accordance with prior trophic classifications (Durkin et al. 2022; Kramer et al. 2025). ASVs assigned to known heterotrophic dinoflagellates and parasitic symbionts (e.g., *Syndiniales* spp.; (Guillou et al. 2008)) were removed (Jones et al. 2025). Remaining dinoflagellate ASVs were assumed to be photosynthetic, though some members of that group (and other phytoplankton groups) are known to be facultatively mixotrophic. Only photosynthetic phytoplankton were included in the resulting dataset.

##### Network-based community detection analysis

A symmetrical adjacency matrix was constructed for all samples based upon the strength of correlations between nodes (here, individual sample sites). Pearson’s correlation coefficients were calculated to describe the edges between nodes for (1) the ratios of the 12 accessory pigments normalized to total chlorophyll-*a* and (2) relative sequence abundances of the 13 18S rRNA gene classes. The edges between nodes were then weighted following Weighted Gene Co-Expression Network Analysis (WGCNA) (Zhang and Horvath 2005): *a*_ij_ = |*corr*(*x_ij_*, *x_i_*)|*^β^*, where *a_ij_* is the adjacency matrix, *corr*(*x_i_*, *x_j_*) is the Pearson correlation coefficient between sites *x_i_* and *x_j_*, and *β* is a scaling term (here *β* = 12 as in (Zhang and Horvath 2005)). The WGCNA was developed to be applied to networks similar to the networks constructed here. These networks have many nodes (324), each of which has many traits, either the ratios of 12 pigments to Tchla or the relative sequences abundances of 13 18S rRNA gene classes (Gates et al. 2016). The “undirected modularity” function was used to perform network-based community detection and separate each sampling site into a distinct community based on its traits (e.g., phytoplankton community composition from pigments or from the 18S rRNA gene) (Rubinov and Sporns 2010). This function forms communities that maximize the modularity of the network, which is a metric that describes network connectivity. Modularity values of 0.3 or greater are considered “high” and indicate robust separation of communities (Fortunato and Hric 2016).

**Figure S1.**
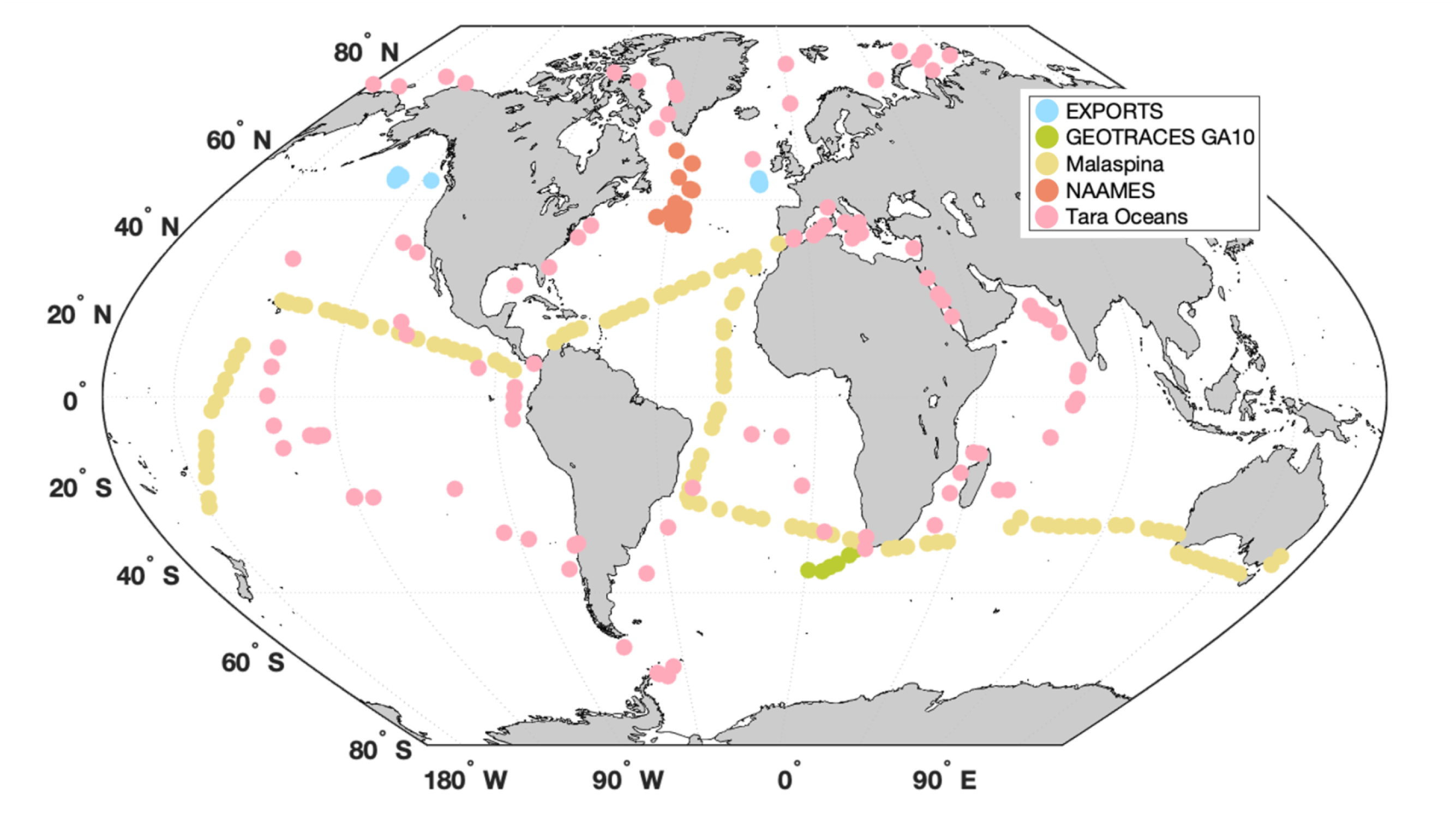
Distribution of samples used in this study colored by cruise (blue = EXPORTS, green = GEOTRACES GA10, yellow = Malaspina, orange = NAAMES, pink = Tara Oceans).

**Figure S2.**
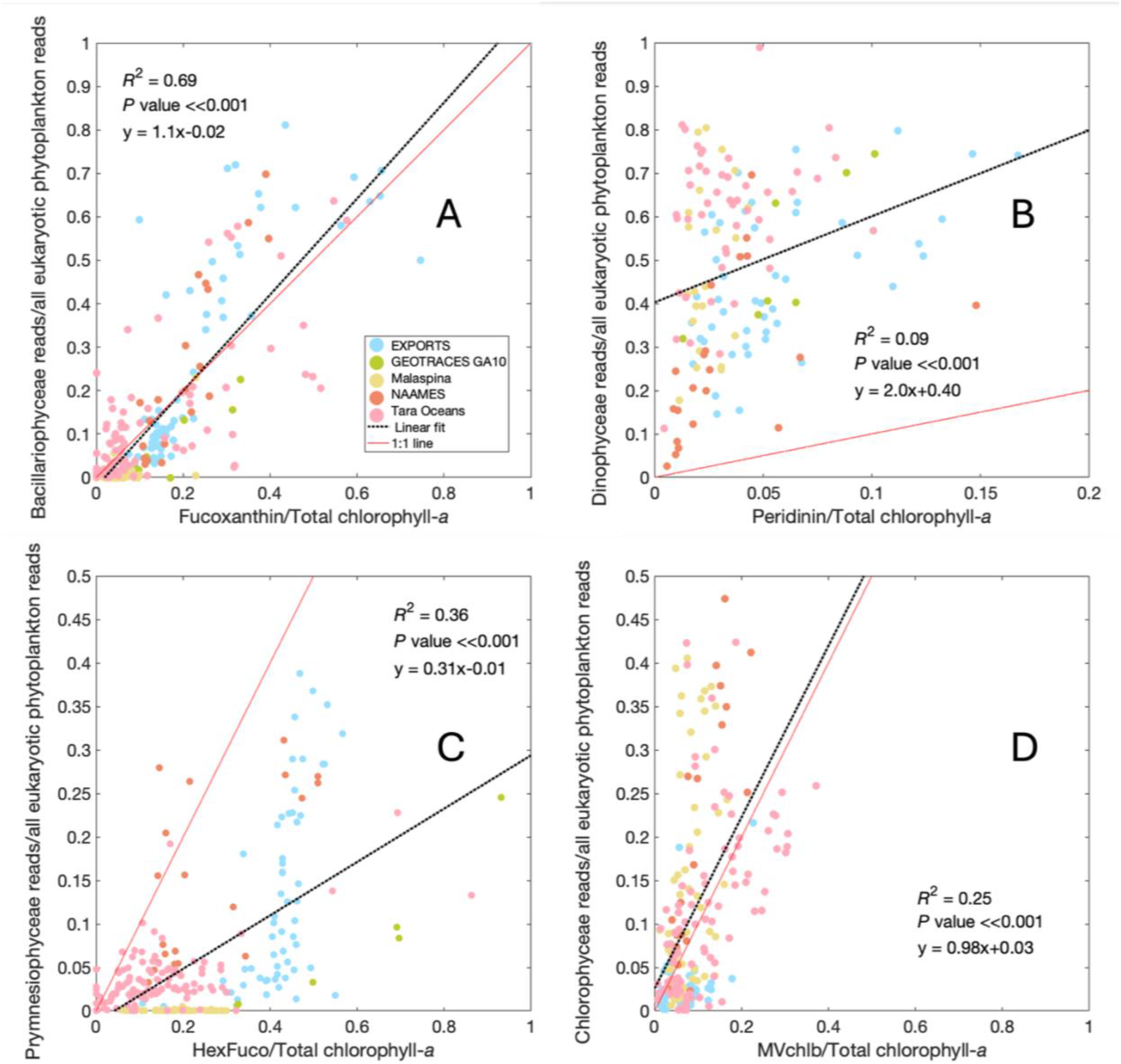
Relationship between (A) diatoms, (B) dinoflagellates, (C) prymnesiophytes, and (D) chlorophytes from 18S rRNA gene sequences (reads / total photosynthetic eukaryotic phytoplankton reads) and from HPLC pigments (pigment / total chlorophyll-*a*). The linear fit is shown for each pairing in a black dashed line and the 1:1 line is a solid red line. Samples are colored by cruise (blue = EXPORTS, green = GEOTRACES GA10, yellow = Malaspina, orange = NAAMES, pink = Tara Oceans).

**Figure S3.**
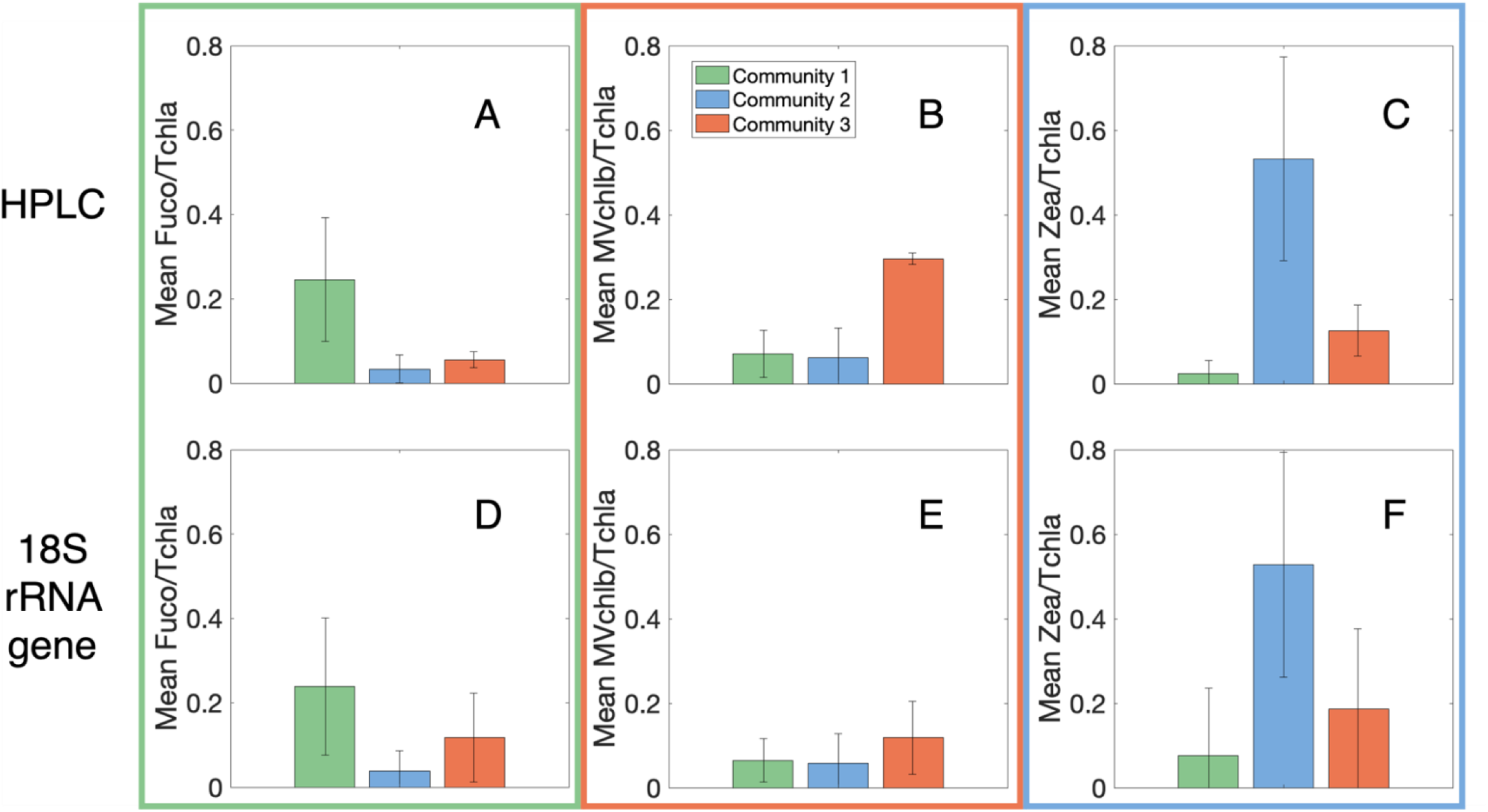
Mean phytoplankton community composition in each community. (A) Mean fucoxanthin (Fuco) to Tchla ratio, (B) mean monovinyl chlorophyll b (MVchlb) to Tchla ratio, and (C) mean zeaxanthin (Zea) to Tchla ratio for the HPLC communities. (D) Mean diatom sequences relative to all phytoplankton sequences, (E) mean Chlorophyte sequences relative to all phytoplankton sequences, and (F) mean Zea to Tchla ratio for the 18S rRNA gene communities. Bars are colored by the community assignment: green = Community 1, light blue = Community 2, orange = Community 3.

**Figure S4.**
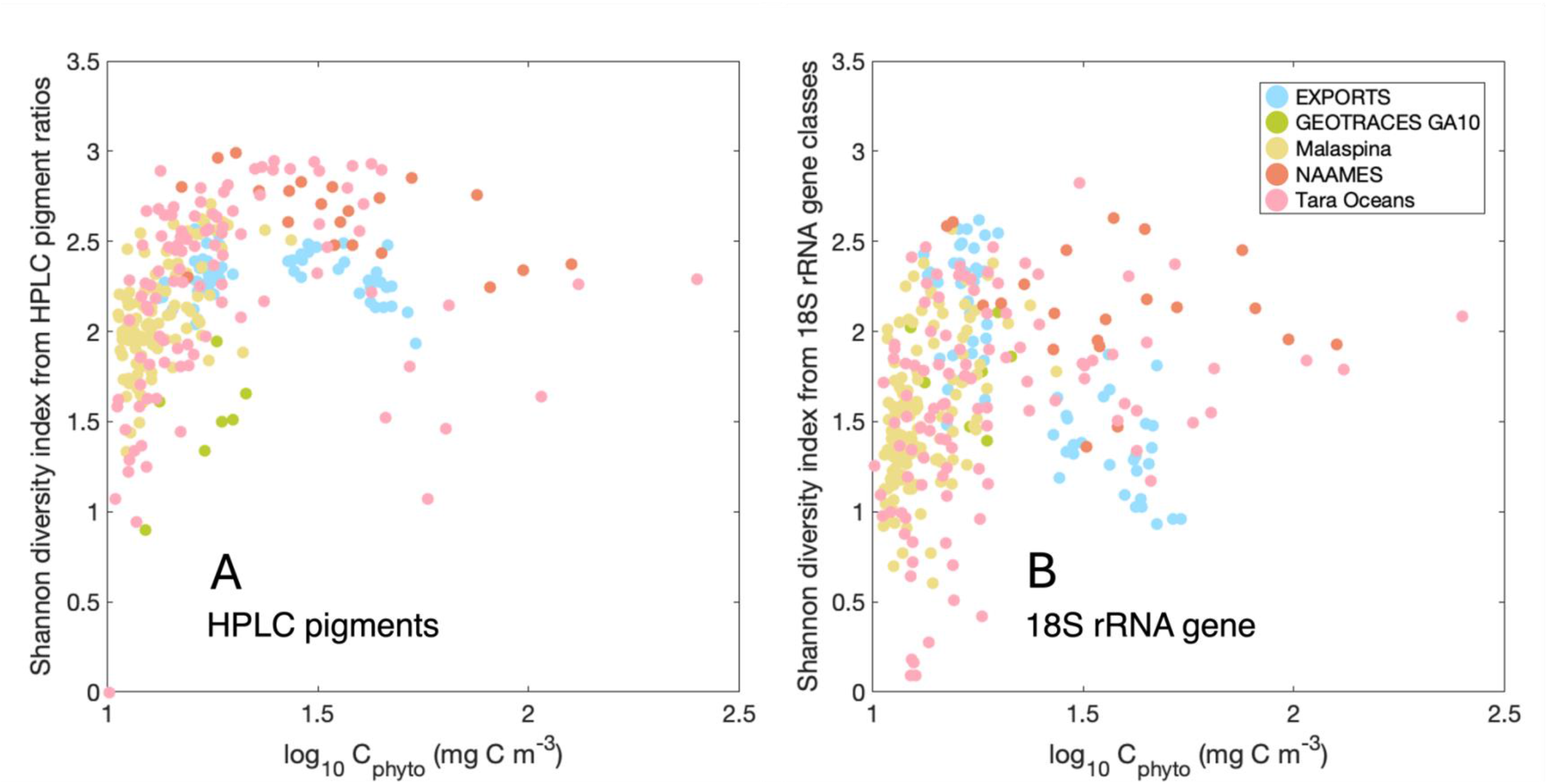
Relationship between phytoplankton diversity based on the Shannon index for (A) ratios of HPLC pigments to total chlorophyll-*a* and (B) relative abundances of 18S rRNA gene classes, with samples colored by cruise (blue = EXPORTS, green = GEOTRACES GA10, yellow = Malaspina, orange = NAAMES, pink = Tara Oceans).

## Notes

https://seabass.gsfc.nasa.gov/experiment/EXPORTS

https://seabass.gsfc.nasa.gov/experiment/NAAMES

https://seabass.gsfc.nasa.gov/experiment/Tara_Oceans_expedition

https://seabass.gsfc.nasa.gov/experiment/Tara_Oceans_Polar_Circle

https://doi.org/10.5281/zenodo.7551643

https://doi.org/10.5281/zenodo.8363876

https://doi.org/10.1594/PANGAEA.948819

https://github.com/jcmcnch/Global-rRNA-Univeral-Metabarcoding-of-Plankton

https://doi.org/10.5285/ff46f034-f47c-05f9-e053-6c86abc0dc7e

https://github.com/sashajane19/Rrs_pigments

### Competing Interest Statement

The authors have declared no competing interest.

### Summary of Updates

1) Clarifying the framing of carbon biomass and phytoplankton diversity throughout to be consistent with the methodological limitations, 2) Revised Figure 2 to reflect nuances in phytoplankton chemotaxonomy, 3) Expanded supplemental figure to show relationships between other major eukaryotic phytoplankton groups observed by both methods (rather than just diatoms), 4) Addition of a short Statistical Analysis section as requested by the journal, and 5) Tightening of the Discussion section to stay focused on the main findings of this manuscript.

